# Metabolic profiling of serum for osteoarthritis biomarkers

**DOI:** 10.1101/2021.12.04.471213

**Authors:** Ziqian Xiao, Zhenyang Zhang, Shanbin Huang, Jerome Rumdon Lon, Shuilin Xie

## Abstract

Osteoarthritis is a prevalent aging disease in the world, and in recent years it has shown a trend toward younger age, which is becoming a major health problem in the world and seriously endangers the health of the elderly. However, the etiology and pathogenesis of osteoarthritis are still unclear, causing great trouble for treatment. To screen out potential biomarkers that could be used as identification of osteoarthritis and explore the pathogenesis of osteoarthritis, we performed untargeted metabolomics analysis of nine New Zealand rabbit serum samples by LC-MS / MS, including three normal serum samples (control group) and six osteoarthritis serum samples (case group). Finally 44 differential metabolites were identified, and the ROC analysis results indicated that a total of 36 differential metabolites could be used as potential biomarkers. Further metabolic pathway enrichment analysis was performed on these differential metabolites, and we found that a total of 17 metabolic pathways were affected, which may provide directions for the study of osteoarthritis mechanisms.

---

Osteoarthritis(OA) is a very common degenerative disease and the incidence rate is increasing with age. In current research, the specific pathogenesis of OA has not yet been investigated. Patients with OA have slow onset in the early stages, with no significant systemic symptoms. It takes up to two years from the onset of pain to the choice to go to hospital, and more than 90% of these patients only go to hospital after they develop knee pain. There are two main types of conventional methods to examine OA, one is imaging, it includes X-ray film examination, irradiation CT, and doing MRI, the other is laboratory tests including hematocrit blood tests, thermal agglutination tests and the examination of the joint fluid in the joint cavity[1]. Imaging is relatively easy, but the probability of misdiagnosis is very high and is often confused with ankylosing spondylitis resulting in medical misdiagnosis. The laboratory inspections are more numerous and complex, and the detection is more difficult.

Considering the large number of biological processes involved in arthritis, a further characterization of the disease mechanisms is needed, which can be used for earlier diagnosis, intervention or treatment. Peng et al. designed a new fluorescence turn-on ADAMTS-4-D-Au probe for detecting ADAMTS-4 activity, which could be used for early diagnosis of cartilage-damage diseases [2]. Leung et al. developed a deep learning prediction model on knee radiographs, which can accurately predicted OA progression in some patients [3]. For the diagnosis of OA, many traditional methods have many defects, and new detection methods are rarely reported, therefore, we focus on metabonomics. Metabolomics is an emerging field that looks for specific metabolic pathways through the study of biomarkers to determine the causes of various diseases from the perspective of metabolite analysis, where lifestyle, diet, disease and genetics can affect multiple metabolite concentrations simultaneously[4]. Some studies in recent years have attempted to identify biomarkers of arthritis through metabolomics and have identified many potential targets in urine, synovial fluid and serum, but specific biomarkers have not yet been identified, so the search for biomarkers through metabolomics is gradually evolving into a new breakthrough direction[5, 6]. For example, before this study, Maerz et al. performed a serum metabolomics analysis after anterior cruciate ligament injury in rats, and conducted a preliminary study on inflammation and immune disorders in traumatic OA[7]. Some metabonomic analyses for OA showed that OA affects the metabolism of amino acids [8–10]. Many biomarkers that could be used as early diagnosis of osteoarthritis have been reported in many studies [10–12]. These corroborate the merits of metabolomic studies for understanding the disease mechanism of arthritis.

The intra-articular injection of papain in rabbits to construct the OA model is a popular method to construct the rabbit knee OA disease model in recent years. In this study, New Zealand rabbits were used as the experimental model, three as the control group, and six were constructed as the experimental group through a noninvasive knee OA modeling method. Papain can degrade chondroitin sulfate (csso4) of cartilage matrix, which increases the amount of free water inside cartilage and affects cartilage elasticity, compressive resistance and integrity of cartilage, so that stresses in the normal range can also cause damage to cartilage, and eventually leading to the occurrence of typical OA[13]. The method of injecting papain to construct OA is to promote disease and will not affect metabonomic analysis. Therefore, we collected rabbit blood samples to prepare serum for metabonomic analysis.

In this project, liquid chromatography tandem mass spectrometry (LC-MS / MS) was used to identify the metabolites and compare the levels of metabolites between the two groups. The differential metabolites were screened by multivariate statistical analysis and univariate analysis, and they were exhibited by clustered heatmap and volcano plot. The diagnostic ability of the screened differential metabolites was judged by ROC analysis to select potential biomarkers for OA. The differential metabolites were finally subjected to pathway enrichment analysis to find out the metabolic pathways affected by OA.

Herein we selected a sample from each group for base peak ion chromatogram (BPC) inspection, the BPC chart is shown in Fig. 1. It shows that the samples have good peak shape and large peak capacity. We performed multivariate statistical analysis and univariate analysis on the processed data by metaX[14], the PCA and PLS-DA[15, 16]score plots are shown in Fig. 2. The PCA analysis shows that the scores of the two principal components in the positive ion mode are PC1=48.48% and PC2=16.04%, and the scores in the negative ion mode are PC1=52.02% and PC2=15.22%. It can be seen that there is an abnormal point in the control group, and this point comes from the sample control-1, indicating that the sample has individual differences. Further PLS-DA[15, 16] analysis was performed on the two groups. Through supervised statistical analysis, the differences between the groups were expanded and the differences within the groups were reduced. From the score plots, it can be seen that the control group is mainly distributed on the right side, while the OA group mainly clustered on the left, the two have a good separation, which indicates that our data can be used for further analysis.

**Fig. 1.**
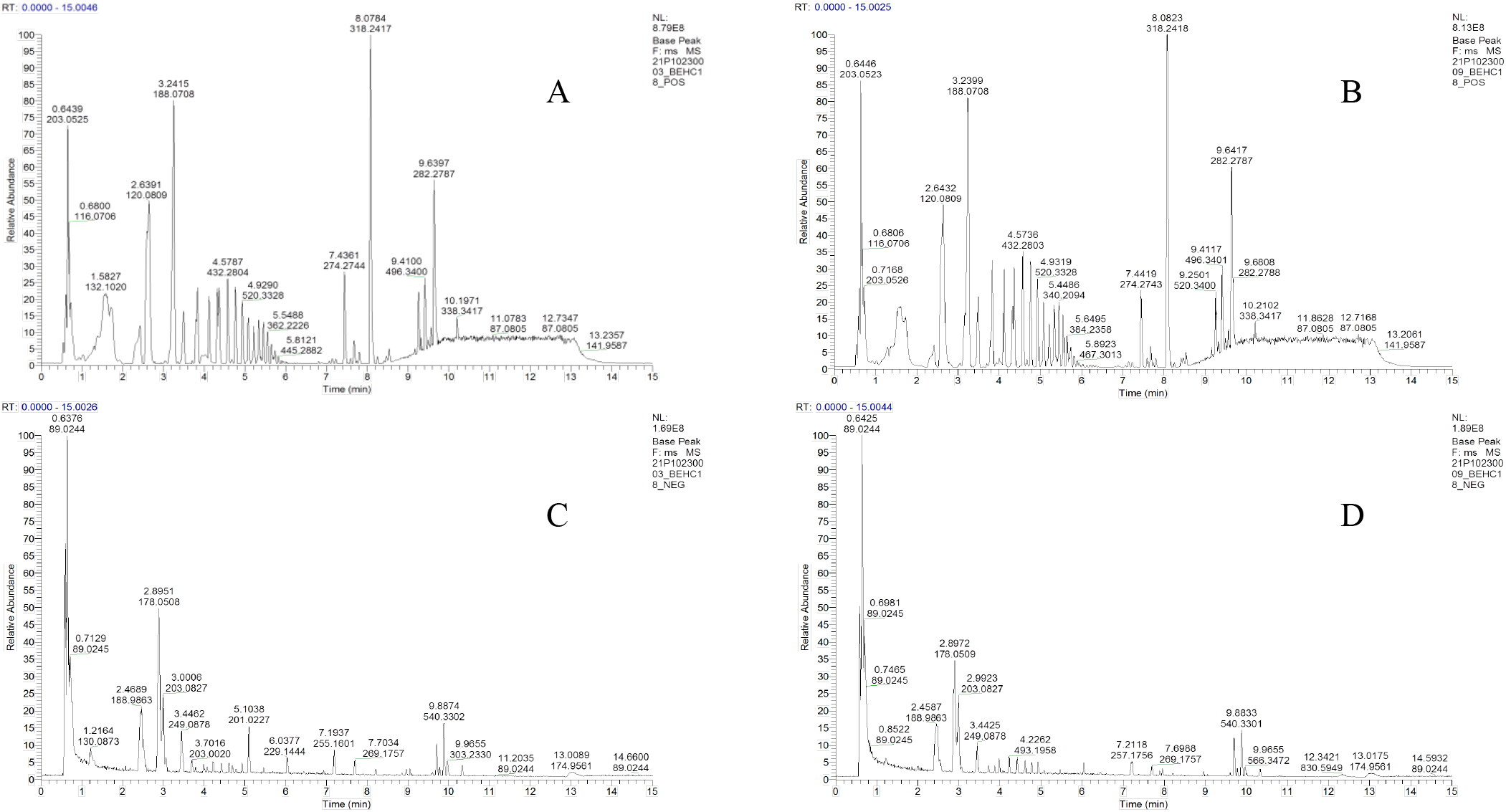
Base peak ion chromatogram (BPC) A. ESI+ BPC-control; B. ESI+ BPC-case; C. ESI-BPC-control; D. ESI-BPC-case

**Fig. 2.**
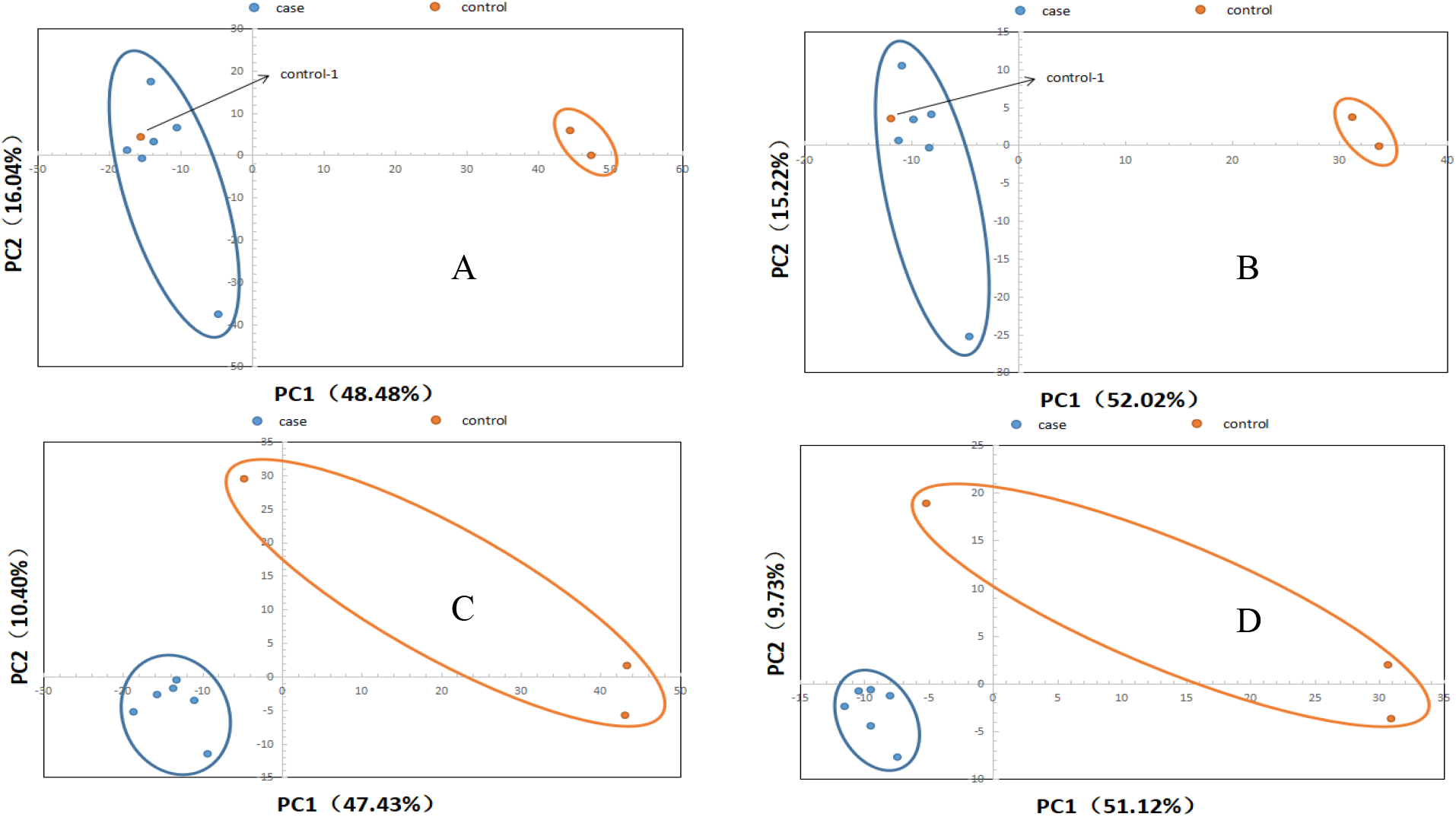
PCA, PLS-DA score plots A. (ESI+) PCA; B. (ESI-) PCA; C. (ESI+) PLS-DA; D. (ESI-) PLS-DA The abscissa is the first principal component PC1, the ordinate is the second principal component PC2. The number is the score of the principal component, which represents the percentage of the explanation on overall variance of the specific pricipal component.

After data preprocessing, 1632 and 636 metabolites were detected in positive and negative ion mode respectively. In order to screen out differential metabolites, fold change analysis and t-test were performed. The screening conditions of differential metabolites were set as fold change ≥ 1.2 or ≤ 0.83, p-value < 0.05. The statistics of screening results are shown in Table 1. The cluster heat maps (Fig. 3) show that all differential metabolites are divided into two clusters of co-regulated metabolites. In the positive ion mode, 33 metabolites in the upper cluster are significantly down regulated and 72 metabolites in the lower cluster are up regulated while in the negative ion mode, 26 metabolites in the upper cluster are up regulated and 8 metabolites in the lower cluster are down regulated. The up-regulated metabolites include fructoselysine, otonecine, carmustine, etc., and the down-regulated metabolites include nepsilon, nepsilon, nepsilon trimethyllysine, tranexamic acid, triethylamine, etc. Draw the differential metabolites into volcano plots (Fig. 4) for visual display, in which red is the up-regulated differential metabolite, green is the down-regulated differential metabolite, and purple-gray is the metabolite with no obvious difference.

**Fig. 3.**
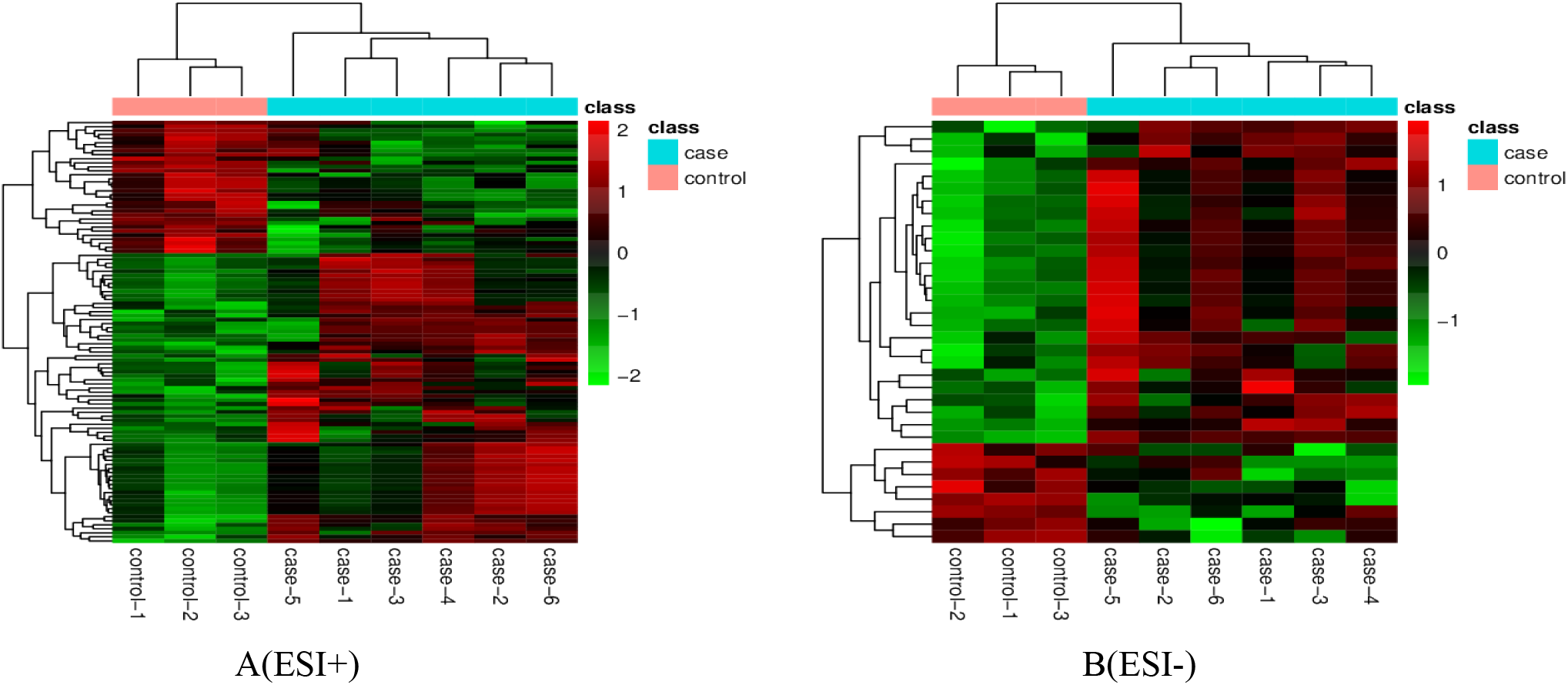
Clustering heat map Each row represents a differential metabolite, each column represents a sample, color is the amount expressed, and green to red indicates the amount expressed from low to high.

**Table 1.**
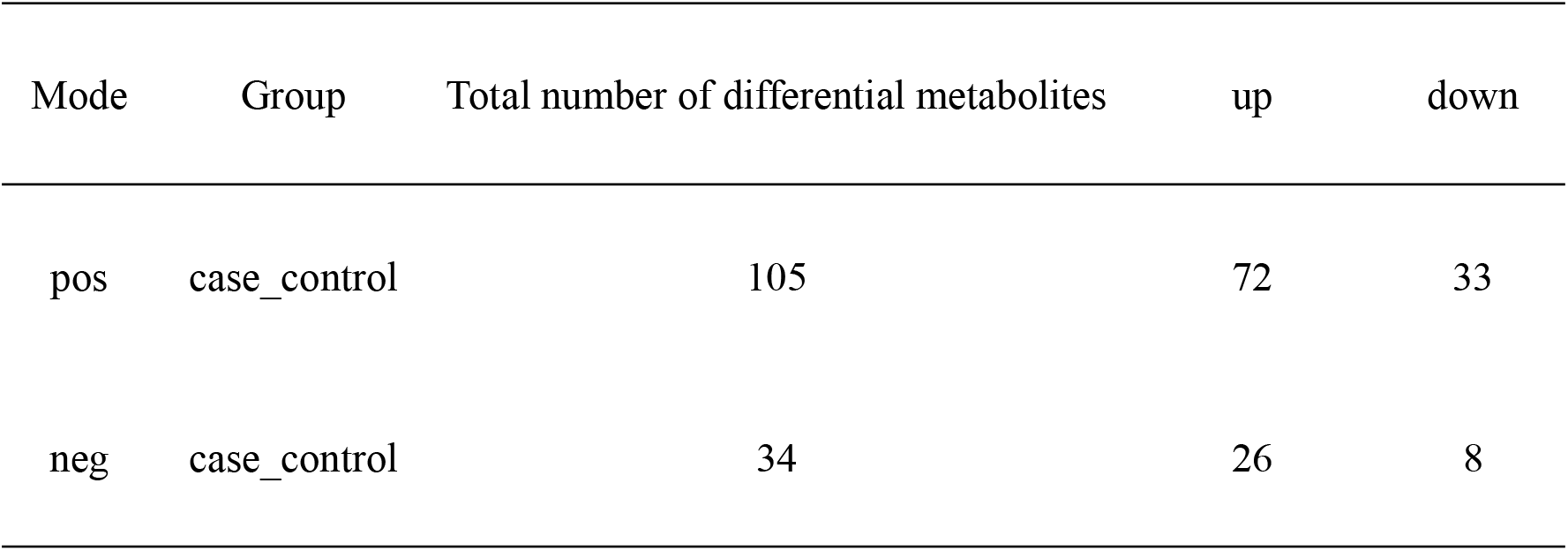
Differential metabolite statistics

| Mode | Group | Total number of differential metabolites | up | down |
| --- | --- | --- | --- | --- |
| pos | case_control | 105 | 72 | 33 |
| neg | case_control | 34 | 26 | 8 |

**Fig. 4.**
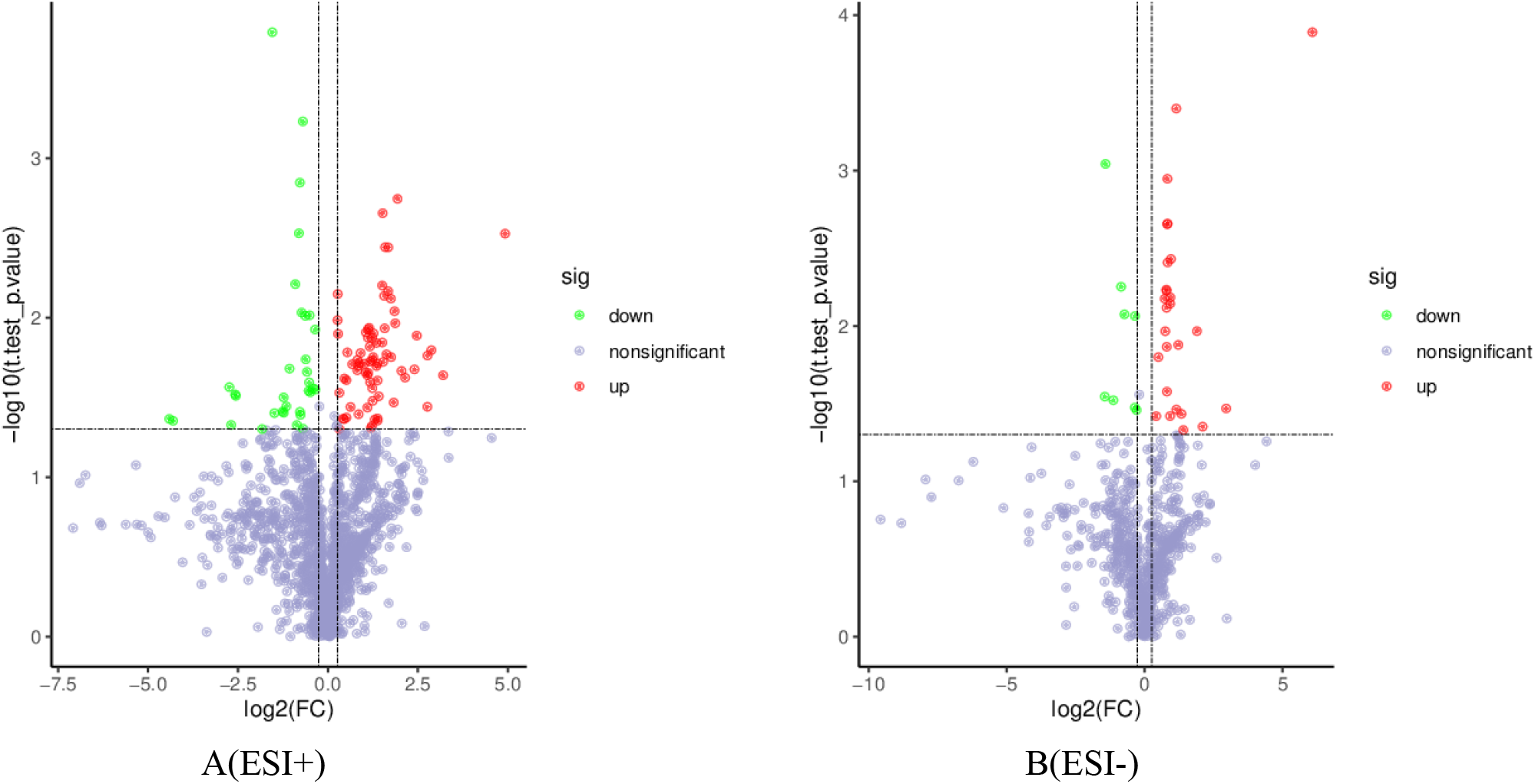
Volcano plots Horizontal axis is log_2_(FC) and vertical axis is −log10 (p-value). The horizontal and vertical dashed lines were selected as the filtering conditions: P < 0.05, log_2_(FC) > 1.2 or < 0.83.The differential metabolites in the top left and the top right part were obtained after filtering, green representing downregulation and red representing upregulation.

We further identified the screened differential metabolites through HMDB database, KEGG database, etc. A total of 35 differential metabolites can be identified in positive ion mode and 9 differential metabolites can be identified in negative ion mode. ROC analysis was performed on these 44 differential metabolites to evaluate their diagnostic ability. The metabolites with area under the curve (AUC) > 0.9 can be used as potential biomarkers. Finally, we determined 36 metabolites as biomarkers of OA, the relevant information is shown in Table 2. These biomarkers include nepsilon, nepsilon, Nepsilon trimethyllysine, ascorbic acid, otonecine, tranexamic acid, etc., all these metabolites have good diagnostic ability for OA.

**Table 2.** List of potential biomarkers

| Name | Fold-change | p.value | direction |
| --- | --- | --- | --- |
| ESI+ |  |  |  |
| Nepsilon,nepsilon,nepsilon-trimethyllysine | 0.3427 | 0.0002 | down |
| Otonecine | 2.2536 | 0.0188 | up |
| Tranexamic acid | 0.0472 | 0.0431 | down |
| Triethylamine | 0.668 | 0.0219 | down |
| Carmustine | 2.8701 | 0.0022 | up |
| Epiguanine | 0.5722 | 0.003 | down |
| Bayer e 39 soluble | 6.8063 | 0.0173 | up |
| Etilevodopa | 9.1913 | 0.023 | up |
| Leu-val | 0.6167 | 0.0006 | down |
| Melatonin | 7.3363 | 0.016 | up |
| 1,4-naphthoquinone | 2.597 | 0.0428 | up |
| Arg-asp | 3.3794 | 0.0177 | up |
| Istamycin a1 | 1.4538 | 0.0165 | up |
| Deferoxamine | 2.3565 | 0.0185 | up |
| Codonocarpine | 1.5971 | 0.0196 | up |
| Sophoranone | 2.3365 | 0.0478 | up |
| Anecortave | 3.1962 | 0.0036 | up |
| Compactin | 2.1724 | 0.0118 | up |
| Quinoline | 0.4259 | 0.0316 | down |
| deoxyphorbol 20-acetate 13-(2-methylbutanoate) | 2.8926 | 0.0189 | up |
| Khellin | 0.6024 | 0.0093 | down |
| Epelsiban | 3.0074 | 0.0036 | up |
| Nevirapine | 0.1551 | 0.047 | down |
| (1r,5r)-3,3,5-trimethylcyclohexyl 5-oxo-l-prolinate | 6.7929 | 0.0362 | up |
| Theaspirane | 0.3567 | 0.0397 | down |
| 2475675 | 0.1698 | 0.0309 | down |
| Drostanolone propionate | 0.1679 | 0.0303 | down |
| ESI- |  |  |  |
| Ascorbic acid | 2.2196 | 0.0004 | up |
| L-(+)-valine | 2.2286 | 0.0345 | up |
| hydroxy-1,2-diphenylethyl hydrogen sulfate | 1.3409 | 0.038 | up |
| 4-carboxy-2-(tyrosylamino)butanoate | 7.7855 | 0.0339 | up |
| Gentisic acid | 0.3758 | 0.0009 | down |
| Benzoic acid | 1.9018 | 0.038 | up |
| (5-amino-2-[(2,6-diamino-2,6-dideoxyhexopyranosyl)oxy]-3-{[3-o-(2,6-diamino-2,6-dideoxyhexopyranosyl)-beta-d-ribofuranosyl]oxy}-4-hydroxycyclohexyl)acetamide | 1.7462 | 0.0136 | up |
| Tryptophyl-d-alanyl-d-allothreonylglycyl-d-histidyl-l-phenylalanyl-d-methioninamide | 2.6554 | 0.0467 | up |
| Probucol | 2.3401 | 0.0132 | up |

The filtered differential metabolites were imported into the KEGG database for pathway enrichment analysis, from which we found that 17 metabolic pathways were affected. Circadian rhythm and vitamin B6 metabolism were detected in positive ion mode, and metabolic pathways detected in negative ion mode included HIF-1 signaling pathway, pantothenate and CoA biosynthesis, mineral absorption, glutathione metabolism etc. The total enrichment results are displayed in the bubble plot for metabolic pathway enrichment analysis (Fig. 5) and the metabolic pathway enrichment results table (Table 3), from which can be seen that the metabolic pathway with the largest enrichment factor is circadian rhythm. Circadian rhythm is a regular cycle established by the adaptation of various physiological functions to changes in the external environment, which also reflects that circadian rhythm has a certain impact on the development of OA. Kc et al.[17] have established circadian rhythm disordered mouse models and found that the disorder of biological clock system lead to pathological changes in the knee joint, suggesting that circadian rhythm disorder will induce the development of OA. Report has also shown that the autonomous clock in chondrocytes regulates key pathways involved in osteoarthritis[18], which can provide more new ideas for the treatment of osteoarthritis by investigating the molecular mechanisms between circadian rhythm and the development of OA. At the same time, it can be seen that the metabolism of multiple amino acids is affected, including valine, leucine and isoleucine biosynthesis and degradation, phenylalanine metabolism, tyrosine metabolism. Meanwhile, up-regulated metabolite L - (+) - valine affects several metabolic pathways, including valine, leucine, and isoleucine biosynthesis, pantothenate and CoA biosynthesis, mineral absorption, and protein digestion and absorption. Many studies have reported the relationship between amino acid metabolism and osteoarthritis[19, 20], and it can be seen that OA has an important impact on the metabolism of relevant amino acids.

**Fig. 5.**
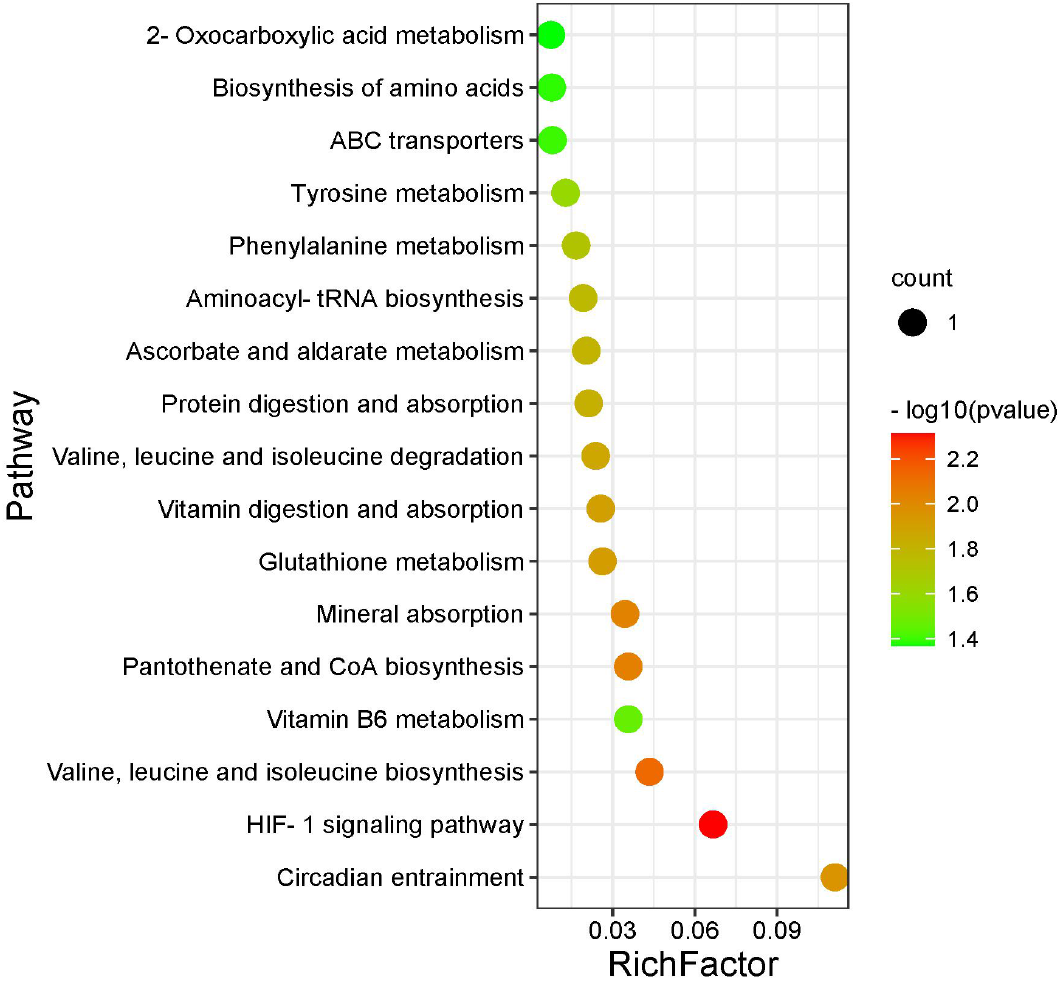
Bubble plots for metabolic pathway enrichment analysis Horizontal axis is enrichment factor and vertical axis is pathway name. Dot size represents the number of differential metabolites annotated to that pathway.

**Table3.** Affected metabolic pathways in rabbit serum

| Pathway | RichFactor | KEGG.Name | KEGG.ID | Pvalue | Direction |
| --- | --- | --- | --- | --- | --- |
| ESI- |  |  |  |  |  |
| HIF-1 signaling pathway | 1/15 | Ascorbic acid | C00072 | 0.0049 | up |
| Valine, leucine and isoleucine biosynthesis | 1/23 | L-(+)-valine | C00183 | 0.0074 | up |
| Pantothenate and CoA biosynthesis | 1/28 | L-(+)-valine | C00183 | 0.0091 | up |
| Mineral absorption | 1/29 | L-(+)-valine | C00183 | 0.0094 | up |
| Glutathione metabolism | 1/38 | Ascorbic acid | C00072 | 0.0123 | up |
| Vitamin digestion and absorption | 1/39 | Ascorbic acid | C00072 | 0.0126 | up |
| Valine, leucine and isoleucine degradation | 1/42 | L-(+)-valine | C00183 | 0.0136 | up |
| Protein digestion and absorption | 1/47 | L-(+)-valine | C00183 | 0.0152 | up |
| Ascorbate and aldarate metabolism | 1/49 | Ascorbic acid | C00072 | 0.0158 | up |
| Aminoacyl-tRNA biosynthesis | 1/52 | L-(+)-valine | C00183 | 0.0168 | up |
| Phenylalanine metabolism | 1/60 | Benzoic acid | C00180 | 0.0193 | up |
| Tyrosine metabolism | 1/78 | Gentisic acid | C00628 | 0.0251 | down |
| ABC transporters | 1/124 | L-(+)-valine | C00183 | 0.0396 | up |
| Biosynthesis of amino acids | 1/128 | L-(+)-valine | C00183 | 0.0408 | up |
| 2-Oxocarboxylic acid metabolism | 1/134 | L-(+)-valine | C00183 | 0.0427 | up |
| ESI+ |  |  |  |  |  |
| Circadian rhythm | 1/9 | Melatonin | C01598 | 0.0111 | up |
| Vitamin B6 metabolism | 1/28 | 4-pyridoxate | C00847 | 0.0343 | down |

Osteoarthritis is a gradually progressive chronic disease, the degenerative damage and reactive hyperplasia of articular cartilage caused by OA are related to many factors, such as increasing age, obesity and strain. OA is often diagnosed by clinical diagnosis and imaging methods, and common imaging methods include radiographs, magnetic resonance imaging (MRI), ultrasound (US), optical coherence tomography (OCT), etc[21]. However, the diagnosis of OA is often not confirmed until late in the disease, whilst approximately half of those identified by imaging methods do not have associated symptoms or disability and the clinical relevance of some radiological features is not completely clear[22, 23]. Biomarkers play an important role in the diagnosis and stage judgment of the disease, and it is important to study the OA related biomarkers in the early diagnosis of OA. Based on this, we hope to screen out the OA related biomarkers and provide a basis for the early diagnosis of OA.

This project performed an untargeted metabolomics analysis of nine New Zealand rabbit serum samples based on LC-MS / MS to extract and compare serum metabolites between control and OA groups. We identified 44 differential metabolites, which were further subjected to ROC analysis, and a total of 36 differential metabolites were screened to serve as potential biomarkers to provide some ideas for the early diagnosis of OA. Through the pathway enrichment analysis of differential metabolites, we found that a total of 17 metabolic pathways were affected, which were worth further study.

The association between OA and circadian rhythms seems to be expected. In fact, a number of studies have shed light on the potential relationship between rheumatoid arthritis and circadian rhythms[24], largely due to the body’s hormone levels vary widely at different times of the day. Circadian rhythms originate from the central pacemaker in the brain’s suprachiasmatic nucleus (SCN) and is regulated by light-sensitive retinal nerves[25]. This rhythm plays an important role in regulating endocrine and immune functions. Some previous studies have shown that melatonin secretion significantly increases in patients with rheumatoid arthritis during the night, while endogenous cortisol synthesis is subsequently activated to counteract the cascade of symptoms[26]. This rhythmic fluctuation may play a role in the pathophysiology of rheumatoid arthritis. Previous studies have found that direct inhibition of pro-inflammatory factors such as melatonin contributes to clinical improvement in rheumatoid arthritis[27]. Some studies revealed that RA risk increases with an unhealthy daily routine[28, 29]. In addition, OA and rheumatoid arthritis are chronic inflammatory diseases. The immune system’s excessive response to OA at night uses up a lot of energy[30, 31]. This disrupts the balance of energy expenditure maintained by the circadian rhythm, which can lead to discomfort beyond the symptoms of arthritis itself[32]. In conclusion, the relationship between chronic inflammation and dysfunctional circadian rhythms in rheumatoid arthritis and OA appears to be complex, with many potential interactions.

Our results showed that OA has great impact on amino acids metabolism, especially on branched chain amino acid (BCAA) metabolism, which had been confirmed by previous reports [8, 10]. Zhai et al. firstly studied the serum-based metabolomics of human OA, and found the ratio of BCAA to histidine has potential clinical use as a biomarker of OA[8]. Although our study did not show changes in histidine content, these studies can reflect the close relationship between BCAA and OA. Study had reported that BCAA is related to the increase number of major pro-inflammatory cytokines involved in the pathophysiology of OA, which can lead to the degradation of articular cartilage matrix [33]. It had also reported that BCAA reflects the relationship between obesity and OA, and obesity is an important factor in the occurrence of OA[34]. In addition, this project showed that valine can be used as a potential biomarker for OA, which had also been reported in previous study [35]. This project also reported some new biomarkers, which have certain significance for the development of new diagnostic methods for OA. In conclusion, the relationship between amino acid metabolism and OA has always been a hot topic.

There are some limitations in this study. OA is easily confused with other types of arthritis such as rheumatoid arthritis, we did not investigate the potential relationship between OA and other arthritic diseases, further validation of whether the screened biomarkers are also able to identify other types of arthritis is needed. Second, the sample size in this study is small, so we could not make further interpretation on the screened biomarkers, meanwhile, we obtained the differential metabolites by database matching, and no further validation was done by comparing with standards, which needs to be done in the future. This study used New Zealand rabbits as the research object, and the genetic information of this species is different from that of human. The future research needs to be closer to the clinical aspect.

## Conclusion

In this project, untargeted metabolomic analysis was performed on nine serum samples by LC-MS / MS. Screening differential metabolites by multivariate statistical analysis combined with the results of fold change analysis and t-test. The screening results were imported into databases for comparison and finally identified 44 differential metabolites. 36 of the 44 metabolites were served as biomarkers for OA by ROC analysis. Pathway enrichment analysis showed that 17 metabolic pathways were affected, including circadian rhythm, vitamin B6 metabolism, HIF-1 signaling pathway, pantothenate and CoA biosynthesis, mineral absorption, glutathione metabolism, etc. This provides new research directions for future therapeutics and development of new drugs in the field of OA.

## Materials and methods

Nine New Zealand rabbit serum samples used for metabolomic analysis were purchased from Guangzhou Huateng Biomedical Technology Co., Ltd. The samples were divided into control group and case group. The control group includes three normal rabbit serum samples named control-1, control-2, control-3, and the case group includes six OA rabbit serum samples named case-1, case-2, case-3, case-4, case-5, case-6. All samples were stored in a refrigerator at − 80 °C.

Serum was stored in −80°C refrigerator before sample preparation. 100 μL samples were extracted by directly adding 300 μL of precooled methanol and acetonitrile (2:1, v/v, Thermo Fisher Scientific, USA). After vortex(QL-901, Kylin-bell Lab Instruments Co., Ltd., China) for 1 min and incubate at −20°C for 2 hours, samples were centrifuged at 14800 rcf for 10 min at 4°C, and the supernatants were then transferred for vacuum freeze drying(Maxi Vacbeta, GENE COMPANY). The metabolites were resuspended in 150 μL of 50% methanol and centrifuged at 14800 rcf for 10 min at 4°C, then the supernatants were transferred to autosampler vials for LC-MS analysis.

Metabolites separation was performed on a Waters 2D UPLC (Waters, USA) with a Waters ACQUITY UPLC BEH C18 column (1.7 μm, 2.1 mm × 100 mm, Waters, USA), and the column temperature was maintained at 45 °C. The mobile phase in positive ion mode is: 0.1% formic acid solution (50144-50ml, DIMKA, USA) (A) - 0.1% formic acid methanol solution (B). The mobile phase in negative ion mode is: 10mM ammonium formate solution (17843-250G, Honeywell Fluka, USA)(A)-10mM ammonium formate in 95% methanol (B). The gradient conditions were as follows: 0-1 min, 2% B; 1-9 min, 2%-98% B; 9-12 min, 98% B; 12-12.1 min, 98% B to 2% B; and 12.1-15min, 2% B. The flow rate was set at 0.35 mL/min and the injection volume was 5 μL.

Primary and secondary mass spectrometry data were collected by a Q Exactive (Thermo Fisher Scientific, USA). The mass spectrometric settings for positive and negative ionization modes were as follows: The full scan range was 70–1050 m/z; spray voltages, 3.80kV/3.20kV; Capillary Temp, 320 °C; Aux Gas Heater Temp, 350 °C; Sheath gas flow rate, 40 arbitrary units; Aux gas flow rate, 10 arbitrary units; runtime, 13 min.

Raw data from mass spectrometry was imported into Compound Discoverer 3.1(Thermo Fisher Scientific, USA) for data processing, including peak extracted, peak alignment, metabolite identification etc., and information about compound molecular weight, retention time, peak area and identification results was exported. Metabolites were identified by using the BGI self-built standard library database, HMDB database, KEGG database, mzCloud database, and Chemspider database. The basis for identification includes Precursor Mass Tolerance< 5 ppm, Fragment Mass Tolerance< 10 ppm, RT Tolerance< 0.2 min. The results exported from the Compound Discoverer 3.1 were imported into metaX[14] for data preprocessing, statistical analysis, and metabolite analysis. Data were normalized using the probabilistic quotient normalization method (PQN[36]) to obtain the relative peak area. Multivariate statistical analysis and univariate analysis were performed on metaX[14]. The distribution and separation trends of the two groups were observed by multivariate statistical analysis, and then fold change (FC) with p-value were obtained by univariate analysis.

Fold change analysis of metabolites was performed between the two groups, t-test was further performed to determine whether the differences in metabolites between groups were significant. Screening for differential metabolites based on the results of univariate analysis, differential metabolite screening conditions were as follows: fold change ≥ 1.2 or ≤ 0.83; p-value < 0.05, and metabolites meeting these conditions could be considered significantly different. For clustering analysis of differential metabolites, data was log2 transformed and Z-score normalized, Hierarchical Clustering was used for clustering algorithms and Euclidean distance was used for distance calculation. Through the cluster analysis, the variation trend of differential metabolites can be seen and find out the metabolites with the same variation trend. After that, the screened metabolites were visual displayed by volcano plots. The filtered differential metabolites were imported into databases for identification, the identified differential metabolites were subjected to ROC analysis, and the differential metabolites with AUC > 0.9 could be tentatively used as biomarkers for OA. The identified differential metabolites were finally imported into KEGG database for metabolic pathway enrichment analysis.

## Acknowledgements

This work was supported by the Science and the Technology Planning Project of Guangzhou (201704020151).

## Author contributions

J.R.L. and S.X. designed the experiments. Z.X. and Z.Z. did the calculation of simulation and analyzed the data. S.H. helped finish the manuscript.

